# Medin Induces Pro-Inflammatory Activation of Human Brain Vascular Smooth Muscle Cells

**DOI:** 10.1101/2024.09.16.613366

**Authors:** Nina Karamanova, Kaleb T. Morrow, Alana Maerivoet, Jillian Madine, Ming Li, Raymond Q. Migrino

**Affiliations:** Phoenix Veterans Affairs Health Care System, Phoenix Arizona USA; University of Liverpool, United Kingdom; University of Arizona College of Medicine-Phoenix, Arizona, USA

**Author notes:** Corresponding Author: Raymond Q. Migrino MD, Phoenix Veterans Affairs Health Care System, 650 E. Indian School Road, Phoenix AZ 85012. Denotes equal first author (Nina Karamanova, Kaleb T. Morrow).

**Keywords:** smooth muscle dysfunction, amyloid, inflammation, aging, vascular dementia, Alzheimer’s disease

## Abstract

**Background:** Medin is one of the most common amyloidogenic proteins and accumulates in the vasculature with aging. Vascular medin accumulation is associated with Alzheimer’s disease, vascular dementia and aortic aneurysms. Medin impairs smooth muscle-dependent vasodilation in isolated human brain cerebral arteries. The role of medin in vascular smooth muscle (VSMC) activation is unknown. We aim to evaluate the effects of medin on human brain VSMC activation.

**Methods:** VSMCs were exposed to physiologic doses of medin (0.5, 1 and 5 µM) without or with small molecule nuclear factor-κB (NFκB) inhibitor RO106-9920 (10 µM) for 20 hours. Polymerase chain reaction, Western blot/enzyme-linked immunosorbent assay were used to quantify gene and protein expressions/secretions, respectively, of pro-inflammatory factors (interleukin (IL)-6, IL-8 and monocyte chemoattractant protein (MCP)-1) and structural and enzyme proteins associated with VSMC phenotypic transformation (smooth muscle actin alpha 2 (ACTA2), myosin heavy chain 11 (MYH11) and NADPH oxidase 4 (NOX4)).

**Results:** Medin exposure increased VSMC gene expression and protein secretion of IL-6, IL-8 and MCP-1 (protein secretion 46.0±12.8x, 20.2±4.1x and 8.7±3.1x, respectively, medin 5 µM versus vehicle, all p<0.05). There was no change in gene or protein expressions of ACTA2, MYH11 and NOX4. Co-treatment with RO106-9920 reduced medin-induced increases in IL-6 and IL-8 and a trend towards reduced MCP-1 secretion.

**Conclusions:** Medin induced pro-inflammatory activation of human brain VSMCs that is mediated, at least in part, by NFκB. Acute medin treatment did not alter structural proteins involved in VSMC phenotypic transformation. The findings support medin as a potential novel mediator of and therapeutic target for vascular aging pathology.

---

Cardio-cerebrovascular diseases (CVD) are the leading causes of mortality and morbidity world-wide and age is the predominant risk factor for these conditions^1^. It is increasingly recognized that vascular aging results in well-defined vascular perturbations independent of the presence of well-established CVD risk factors such as hypertension, diabetes, hyperlipidemia and smoking^2^. The classic phenotypic changes associated with vascular aging include endothelial and smooth muscle dysfunction, vascular inflammation, calcification and stiffening and prothrombotic transformation^2^. The molecular drivers of vascular aging-induced pathologies remain poorly understood. Medin is a 50 amino acid cleavage product of milk fat globule-EGF factor 8 protein (MFGE8) and forms one of the most common yet least studied human amyloidosis^3^. Medin has been implicated in the pathophysiology of aortic aneurysms^4-7^, vascular dementia^8,9^ and Alzheimer’s disease(AD) ^8,10^, including recent preclinical discovery of its role in aging-related cerebrovascular dysfunction^11^ and co-aggregation with β-amyloid in cerebral amyloid angiopathy^10^ in aged and AD mouse models, respectively. Medin was shown to impair endothelium and smooth-muscle dependent vasodilation in human cerebral and adipose arterioles through oxidative and nitrative stress^12,13^ and medin induced human endothelial cell pro-inflammatory activation via nuclear factor-κB (NFκB) activation^9,13,14^. Medin amyloid accumulates predominantly in the medial layer of arteries^3^ and vascular smooth muscle cells (VSMC) express the highest amount of MFGE8 among various cell types, second only to cone photoreceptor cells^15^. Medin exposure resulted in oxidative stress, reduced VSMC viability^7^ and increased production of matrix metalloproteinase-2^16^. We hypothesize that medin contributes to vascular aging phenotype by causing VSMC pro-inflammatory activation and phenotypic switching from a functional contractile state to an activated synthetic/proliferative state. The aims of the study are to measure the effects of medin exposure in human brain VSMCs on gene and protein expression of pro-inflammatory cytokines and structural proteins associated with VSMC phenotypic switching.

## Methods

The data that support the findings in this study are available upon reasonable request from the corresponding author.

### Recombinant Medin Production

Lemo 21 (DE3) cells were used to express medin utilizing pOPINS-medin with the detailed description of preparation and purification described in prior work^12^. Purity of medin was confirmed to be >95% by SDS-PAGE, measured by matrix-assisted laser desorption and ionization mass spectrometry. *Limulus* Amebocyte Lysate assay (Pierce, Dallas TX) was used to confirm that endotoxin levels were <0.5 ng/mL.

### Human Brain VSMC and 3T3 cell treatments

Primary culture human brain VSMCs (passages 4-8; ScienCell Research Laboratories, Carlsbad CA) were seeded into 6 well plates and grown to full confluence. VSMCs were exposed to 0, 0.5, 1 and 5 µM medin for 20 hours. The medin doses selected are consistent with levels seen in human tissues^13^. Cell lysates were collected for measurement of gene expression of interleukin (IL)-6, IL-8, monocyte chemoattractant protein (MCP)-1, smooth muscle actin alpha 2 (ACTA2), myosin heavy chain 11 (MYH11) and NADPH oxidase 4 (NOX4). Western blot for protein expression of ACTA2 and NOX4. IL-6, IL-8 and MCP-1 are pro-inflammatory cytokines generated by activated VSMCs^17,18^. ACTA2 and MYH11 are contractile proteins whose expressions are downregulated during VSMC proliferative transformation^19^. VSMCs switch from differentiated to less differentiated phenotype during activated proliferative state and this change was shown to be marked by reduced NOX4 expression^20^. Gene expression was measured using PCR (all primers obtained from IDT DNA Technologies, Coralville IL) with β-actin as reference normalization gene, similar to previously reported methods^9^. Primers used were: (IL)-6 (5’-AAC CTG AAC CTT CCA AAG ATG -3’ F and 5’-TCT GGC TTG TTC CTC ACT ACT - 3’ R), IL-8 (PrimeTime qPCR Primers, Exon Location: 1-1, RefSeqNumber:NM_000584), monocyte chemoattractant protein (MCP)-1 (Primer 1: 5’-TCT TTG TCT TCT CCT GCC TGC CTT, Primer 2: 5’-TTA TGA GGC TTG TCC CTT GCT CCA), smooth muscle actin alpha 2 (ACTA2) (PrimeTimeqPCR Primers, Exon location:7-8, RefSeqNumber: NM_001613), myosin heavy chain 11 (MYH11) (PrimeTimeqPCR Primers, Exon location:7-8, RefSeqNumber: NM_022844) and NADPH oxidase 4 (NOX4) (PrimeTimeqPCR Primers, Exon location:1-2, RefSeqNumber: NM_0139550). Western blot was performed using antibodies to ACTA2 (mouse monoclonal, clone 1A4 Sigma-Aldrich, St. Louis MO, dilution 1:4000) and NOX4 (rabbit polyclonal NB110-58851, Novus Bio, Centennial CO, dilution 1:500). IL-6 and MCP-1 in conditioned media were measured with enzyme linked immunosorbent assay (ELISA) using Bio-Rad Bio-Plex Human Cytokine Screening Panel Express and measured in the Bio-Rad Bio-Plex 200 System (Bio-Rad Laboratories, Hercules CA). Secreted IL-8 was measured from conditioned media using IL-8 DuoSet ELISA (R&D Systems, Minneapolis MN).

We previously showed that endothelial cell pro-inflammatory activation by medin, manifested by increased IL-6, IL-8 and IL-1b, was suppressed by RO106-9920, a small molecule specific inhibitor of NFκB^9^. To directly measure the effect of medin on NFκB activation, we utilized an available NFκB luciferase reporter in 3T3 mouse fibroblast cells (Signosis, Inc., Santa Clara CA). VSMCs, endothelial cells and fibroblasts share common response of NFκB activation with oxidative stress^21^ and it was reported that the differences between activated fibroblasts and phenotypically modulated VSMCs may be very subtle or nonexistent^22^. 3T3 cells were exposed to vehicle or medin (5 µM) for 1, 6 or 20 hours and luminescence was measured following addition of luciferase substrate using luminometer (GloMax Navigator System, Promega, Madison WI).

In separate experiments to assess the role of NFκB in pro-inflammatory activation, we utilized RO106-9920, a small molecule specific inhibitor of NFκB^13,23^. VSMCs were treated for 20 hours and 3T3 cells were treated for 6 hours with the following: vehicle, medin 5 µM without and with RO106-9920 (10 µM) or RO106-9920 (10 µM) alone. For VSMCs, IL-6, IL-8 and MCP-1 gene expressions were measured using PCR in cell lysates while their secretion was measured by ELISA of conditioned media. For 3T3 cells, luminescence was measured using luminometer.

### Data and Statistical Analyses

Data are expressed as mean±standard error of means. Significant p value was set at p<0.05 (two-sided). Each data point (except for data in Figure 2A) is a biologic replicate of independent experiments consistent with National Institutes of Health Rigor and Reproducibility Standards. Data that do not have normal distribution underwent log normal transformation and the normally distributed transformed values were used for statistical analyses. Paired group analyses were done using one-way repeated-measures analysis of variance with Tukey’s post hoc pairwise testing using GraphPad Prism 9 (Graphpad Software Inc., San Diego CA).

## Results

VSMCs treated for 20 hours with medin showed increased gene expression of IL-6, IL-8 and MCP-1 at all 3 doses used (Figure 1A-C). Although there was a trend towards increasing response with increasing doses, the differences among the medin doses were not statistically significant. On the other hand, there was no difference in gene expression of ACTA2, MYH11 and NOX4 between medin and vehicle treated cells (Figure 1D-F). Consistent with this finding, there was also no change in protein expression of ACTA2 and NOX4 following medin treatment (Figure 1G-H).

**Figure 1.**
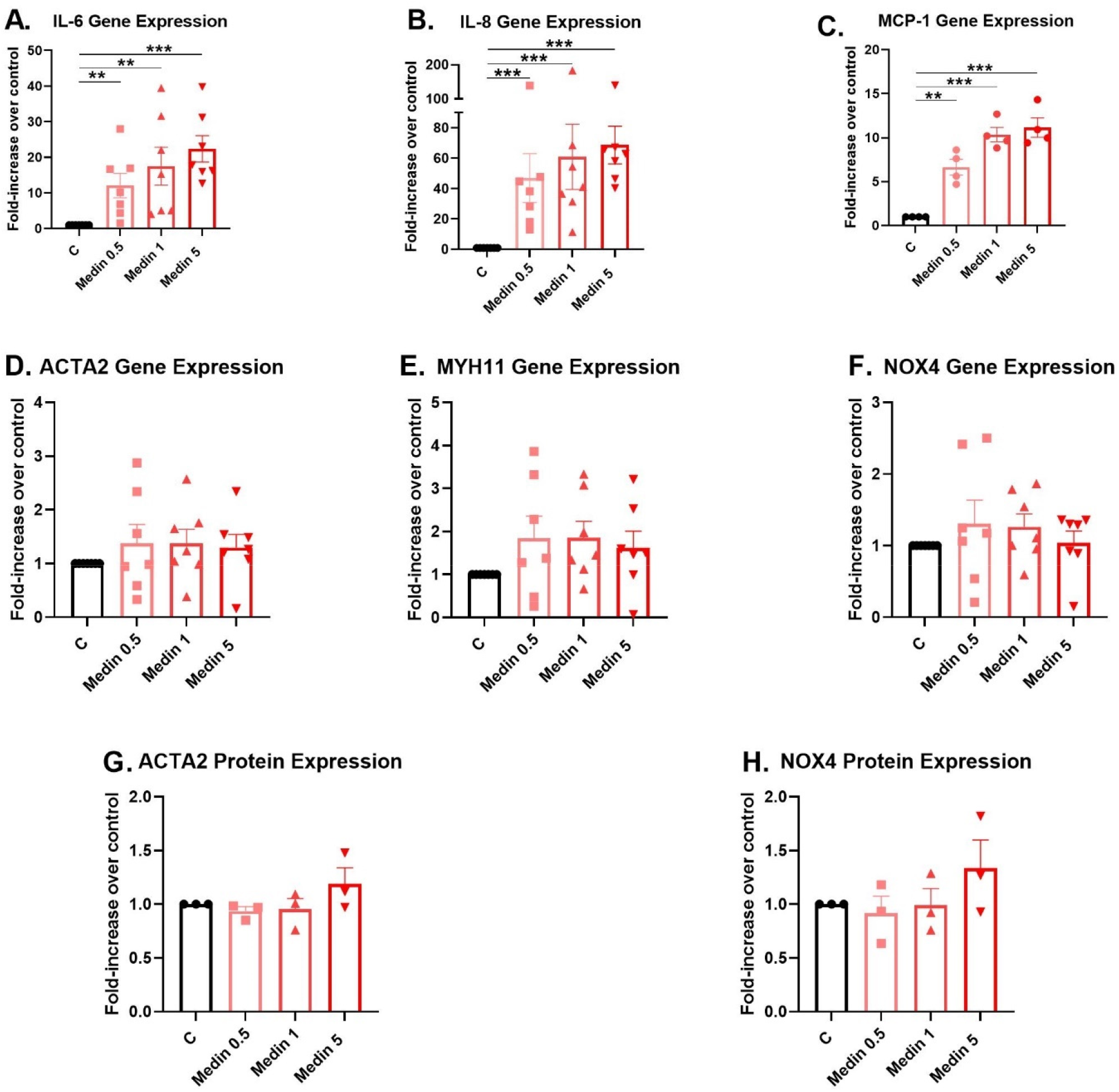
Medin effects on VSMC pro-inflammatory cytokine and structural protein production. A-C. VSMCs treated with medin for 20 hours showed increased gene expressions of IL-6, IL-8 and MCP-1. Although there were progressive increases in levels with higher medin doses, the differences among medin doses were not significant. D-H. Medin did not alter VSMC gene expression of ACTA2, MYH11 and NOX4 and protein expression of ACTA2 and NOX4. **p<0.01, ***p<0.001

3T3 cells with NFκB luciferase showed increased NFκB activation 1 hour following medin treatment, further increasing by 6 hours and peak levels reduced by 20 hours, with levels remaining higher than vehicle control at all time points (Figure 2A). Medin induced NFκB activation in all 3 treatment doses at 6 hours (Figure 2B). Co-treatment with RO106-9920 partially reversed medin’s effect on 3T3 NFκB activation (Figure 2C).

**Figure 2.**
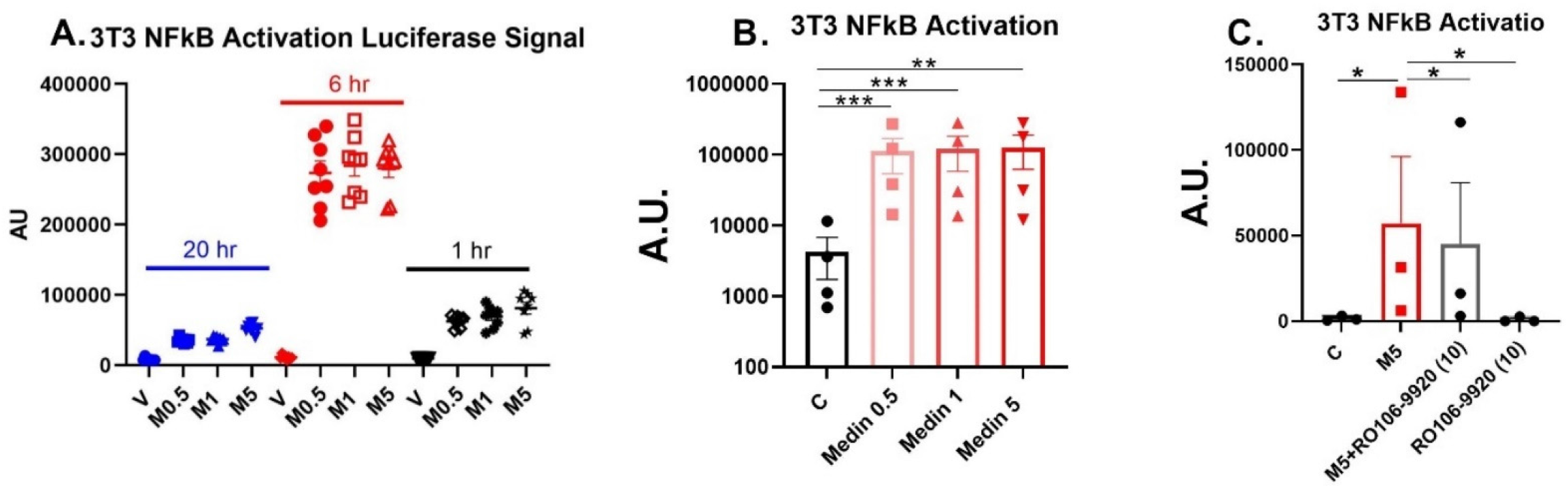
Medin effects on NFκB activation. A. 3T3 cells exposed to medin (5 µM) showed increased luminescence at 1, 6 and 20 hours versus vehicle control with highest luminescence noted at 6 hours. (N=1 biologic replicate, each datapoint is a technical replicate). B-C. 3T3 cells exposed to medin for 6 hours showed increased luminescence that was partially reversed by RO106-9920. *p<0.05, **p<0.01, ***p<0.001

Co-treatment of medin with RO106-9920 in VSMCs showed a trend (not statistically significant) towards reduced IL-6, IL-8 and MCP-1 gene expression versus medin treatment (Figure 3A-C). On the other hand, co-treatment with RO106-9920 showed partial reversal of medin-induced increases in IL-6 and IL-8 protein secretion and a trend towards reduced MCP-1 protein secretion (Figure 3D-F).

**Figure 3.**
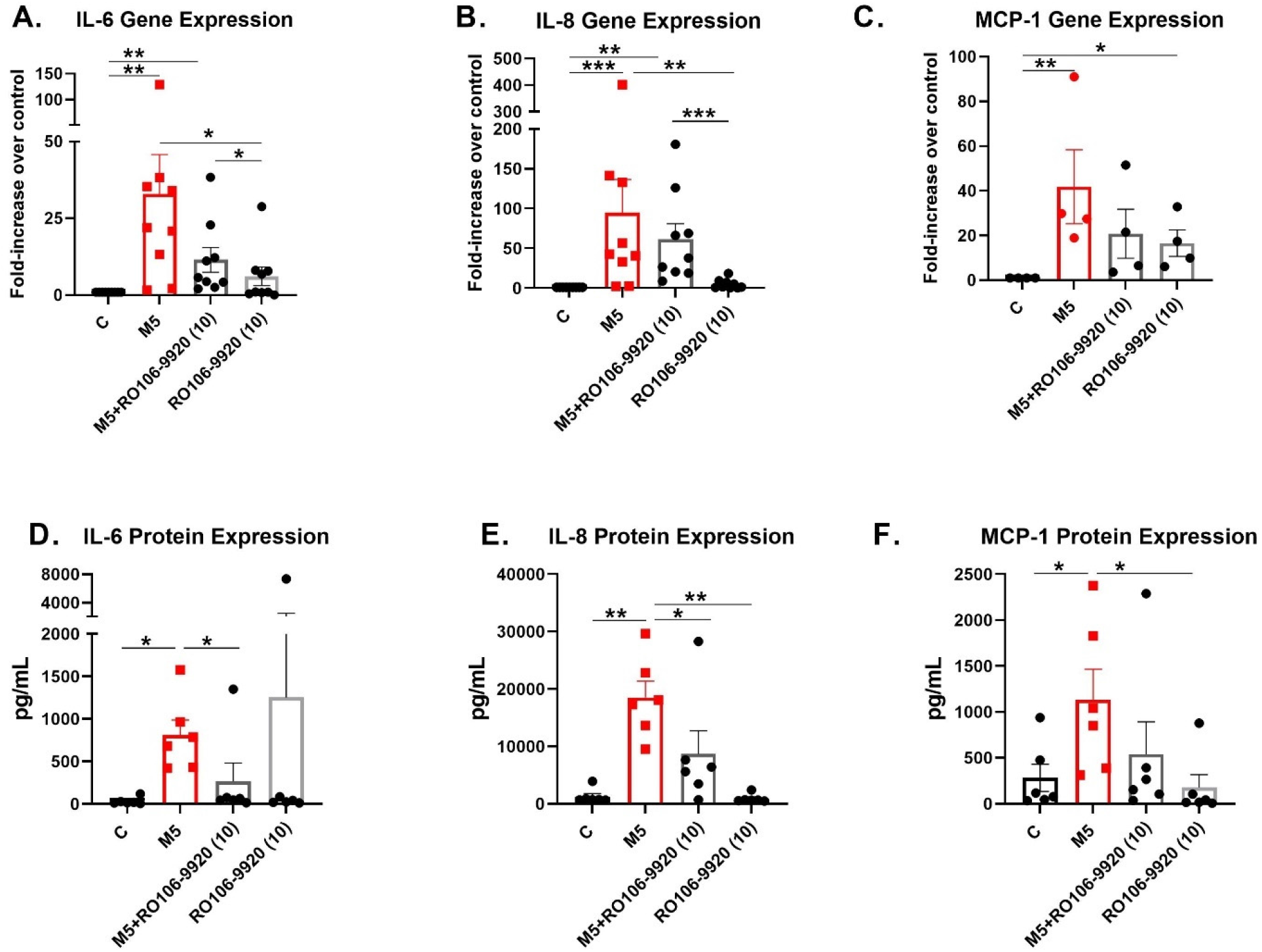
Medin-induced pro-inflammatory activation and NFκB inhibitor. VSMCs treated with medin 5 µM showed increased gene expression (A-C) and protein secretion (D-F) of IL-6, IL-8 and MCP-1. Co-treatment of medin with RO106-9920 attenuated the increased protein secretion of IL-6 and IL-8 and a trend towards reduction of MCP-1. *p<0.05, **p<0.01, ***p<0.001

## Discussion

VSMCs play an important role in arterial aging pathologies as aging-associated VSMC pro-inflammatory activation and phenotypic changes lead to vascular degeneration, extracellular matrix degradation and alterations in mechanical properties leading to arterial stiffness^24^. Medin was originally discovered as the protein component of aortic medial amyloid and it is considered the most common human amyloid as it accumulates with aging throughout the vasculature, including cerebral, coronary, peripheral arteries and aorta^3,25^. Little is known about the effects of medin on VSMCs, the dominant and most abundant cell type in the media layer of arteries. VSMCs have one of the highest expressions of MFGE8^15^ and are therefore likely to be a major source of medin production. VSMCs play a key role in cell-to-cell crosstalk with endothelial cells and immune cells during inflammation^17^. In terms of medin’s effect on VSMC function, we previously showed that rapid autopsy-derived human cerebral arteries from cognitively normal, AD or vascular dementia donors had impaired smooth muscle-dependent vasodilation when treated with physiologic dose of medin for 1 hour^12^. Our results show that medin induces a substantial pro-inflammatory activation in VSMCs with 46.0±12.8x, 20.2±4.1x and 8.7±3.1x increased protein secretion of IL-6, IL-8 and MCP-1 versus vehicle control, respectively (using 5 µM medin). Prior data in endothelial cells and additional new data in 3T3 cells show that medin induces NFκB activation; co-treatment of medin 5 µM with NFκB inhibitor RO106-9920 partially (using 10 µM dose) or fully (using 100 µM dose) reversed increased IL-6 or IL-8 in endothelial cells induced by medin^13^. Our data also shows partial reversal of medin-induced increases in IL-6 and IL-8 (and trend towards similar response for MCP-1) with RO106-9920 suggesting that NFκB activation is also, at least in part, contributing to the pro-inflammatory effect of medin in VSMCs. VSMCs did not tolerate 100 µM of RO106-9920 and it is possible that doses higher than 10 µM could rescue the effects even further.

Our results are significant in terms of understanding potential etiopathology of vascular aging and atherosclerosis. IL-6, IL-8 and MCP-1 are potent mediators of inflammation that contribute to atherogenesis and these cytokines contribute to the synergistic interaction between VSMCs and monocytes^26^. Moreover, medin-induced brain VSMC inflammatory activation may have implications for neuroinflammation. Medin amyloid was found to be more abundant in the media and intima layers of cerebral arteries of vascular dementia and AD donors when compared to cognitively normal brain donors^8^. Evidence suggests that vascular inflammation could initiate or modulate neuroinflammation. In vascular dementia, activated astrocytes and microglia (cellular markers or neuroinflammation), are commonly found concentrating around blood vessels^9,27,28^ and it was shown that medin-induced pro-inflammatory activation of endothelial cells could augment astrocyte activation^9^. There is support for the notion that neuroinflammation is not just an epiphenomenon but may directly contribute to neurodegeneration and cognitive dysfunction^27,29^.

Vascular disease is associated with VSMC plasticity-dependent transformation from a contractile differentiated to a synthetic de-differentiated phenotype^17^. This transformation is usually associated with changes in structural protein and enzyme markers with reduced expression of ACTA2, MYH11 and NOX4^17,19,20^. Our results show no change in gene or protein expression of these proteins in medin-treated VSMCs. This suggests that although medin induces a profound pro-inflammatory activation of VSMCs, acute treatment does not lead to transformation to a de-differentiated state. Note that the effects of chronic medin exposure on VSMC transformation remain unknown.

### Limitations

The study has important limitations. The study was limited to acute treatment and effects of chronic exposure, similar to *in vivo* conditions were not explored. This issue may be more relevant as to the chronic effects of medin on VSMC phenotypic transformation. Inflammatory cytokine activation transcription pathways not dependent on NFκB such as activator protein-1, activating transcription factors and cAMP-response element binding protein were not explored in the current study and should be pursued in future studies. Mediators upstream of NFκB need to be explored; previously we showed that medin induces oxidative and nitrative stress in endothelial cells^12,13^ and oxidative stress has been shown previously to activate NFκB in endothelial cells, smooth muscle cells and fibroblasts^21^.

## Conclusions

Medin induced pro-inflammatory activation of human brain VSMCs that is mediated, at least in part, by NFκB. Acute medin treatment did not alter structural proteins involved in VSMC phenotypic transformation. The findings support medin as a potential novel mediator of and therapeutic target for vascular aging.

## Acknowledgments

We thank Ms. Gail Farrell, the Arizona Veterans Research and Education Foundation and Phoenix VA Office of Research for their support. The views expressed here do not represent the views of the US Department of Veterans Affairs or the United States government.

